# Copper deficiency drives OXPHOS impairment and mitochondrial hyperfusion via MTCH2 in skeletal muscle

**DOI:** 10.1101/2025.11.19.688750

**Authors:** Young-Seung Lee, Hee Soo Kim, Phuc L. Nguyen, Junhyeong Lee, Dong-Il Kim, Jeongmin Lee, Changjong Moon, Kyoung-Oh Cho, Byung-Eun Kim, Jiyun Ahn, Timothy F. Osborne, Taner Duysak, Jeong-Sun Kim, Chang Hwa Jung, Tae-Il Jeon

## Abstract

Copper is an essential trace element for mitochondrial respiration and cellular metabolism, yet its physiological role in skeletal muscle remains incompletely understood. Here, we show that skeletal muscle-specific deletion of the high-affinity copper importer *Ctr1* (SMKO) in mice causes local copper deficiency, resulting in exercise intolerance, systemic metabolic dysfunction, and hallmarks of mitochondrial myopathy such as ragged-red fibers, lactic acidosis, and aberrant mitochondrial morphology. Mechanistically, copper starvation disrupted the electron transport chain proteome and drove pathological mitochondrial hyperfusion. We identified mitochondrial carrier homolog 2 (MTCH2), an outer mitochondrial membrane protein, as a copper-binding regulator that coordinates mitochondrial copper distribution and morphology. Restoring copper levels via a copper ionophore or AAV-mediated *Ctr1* re-expression rescued mitochondrial function and alleviated myopathic features in SMKO. These findings uncover the functional coupling of CTR1 and MTCH2 as a critical mechanistic link between copper homeostasis and mitochondrial remodeling required for skeletal muscle function.

## INTRODUCTION

Copper is an essential trace element that underpins a vast array of physiological processes. In addition to serving as an indispensable cofactor for metabolic enzymes, copper directly modulates signaling pathways that dictate cell growth, differentiation, and survival.^1,2^ Because both copper deficiency and overload can impair these critical functions and drive human disease, systemic and intracellular copper levels are subjected to tight regulation.^3^ In mammals, dietary copper is absorbed in the intestine and exported into the bloodstream by ATP7A copper-transporting ATPase.^4^ Circulating copper is subsequently imported into cells via the high-affinity copper transporter 1 (CTR1) and distributed by intracellular copper chaperones like antioxidant 1 (ATOX1), which delivers copper to the trans-Golgi network to support cuproenzyme biosynthesis.^5^ To prevent toxicity under conditions of copper excess, export machinery is rapidly mobilized to the plasma membrane or vesicular compartments.^6,7^

As the only known copper-specific importer in mammals, CTR1 is broadly expressed and localized to the plasma membrane, where it facilitates constitutive copper uptake under physiological conditions.^5^ The necessity of this importer is highlighted by genetic models: intestinal-specific deletion of *Ctr1* in mice causes perinatal lethality due to systemic copper deficiency,^8^ while cardiac-specific knockout results in progressive cardiomyopathy and premature death.^9^ These models underscore the non-redundant requirement for CTR1 in maintaining tissue-specific copper homeostasis.

Skeletal muscle comprises approximately 40% of total body mass and is a primary driver of systemic energy metabolism.^10,11^ Muscle performance heavily relies on distinct myofiber subtypes–mitochondria-dense oxidative (slow-twitch) fibers that sustain endurance through oxidative phosphorylation (OXPHOS), and glycolytic (fast-twitch) fibers that power short bursts of activity.^12,13^ Consequently, mitochondrial fitness is a critical determinant of muscle health, and its impairment is central to diverse myopathies.^14–17^ Within the mitochondria, copper is obligatorily required for the assembly and activity of cytochrome c oxidase (COX, Complex IV) to sustain the electron transport chain (ETC).^18,19^ Furthermore, copper deficiency is associated with striking alteration in mitochondrial morphology, including the upregulation of fusion proteins such as mitofusin (MFN) 2 and optic atrophy protein 1 (OPA1); however, the precise molecular mechanisms driving this architectural remodeling remain elusive.^20^

While skeletal muscle harbors approximately 27% of total body copper–and muscle weakness is a hallmark of inherited copper transport disorders like Menkes disease^21,22^–the physiological role of copper in mature skeletal muscle remains poorly understood. *In vitro* studies indicate that both cellular copper import via CTR1 and mitochondrial uptake via the inner membrane transporter SLC25A3 are required for myogenic differentiation and COX activity.^23,24^ However, whether copper uptake is essential for skeletal muscle function *in vivo*, particularly through the regulation of mitochondrial dynamics, remains unknown.

Here, we show that skeletal muscle-specific *Ctr1* knockout (SMKO) mice exhibit severe intracellular copper depletion, which drives impaired OXPHOS, a reduction in type IIB muscle fibers, progressive myopathy, and mitochondrial hyperfusion. Mechanistically, we identify mitochondrial carrier homolog 2 (MTCH2), an outer mitochondrial membrane (OMM) protein, as a dual-function, copper-binding coordinator of both mitochondrial copper trafficking and morphological dynamics. Restoring intracellular copper via the ionophore elesclomol (ES) or adeno-associated virus (AAV)-mediated *Ctr1* re-expression fully rescues mitochondrial function and maintains muscle integrity in SMKO mice. Together, these findings reveal an absolute requirement for copper in skeletal muscle maintenance and uncover how MTCH2-mediated copper sensing translates cellular copper dyshomeostasis into mitochondrial architectural remodeling.

## RESULTS

### Exercise-induced copper uptake is required for skeletal muscle performance

To investigate the relationship between trace metal dynamics and physical exertion, we quantified metal levels in the gastrocnemius (GA) muscle of wild-type mice following a four-week treadmill training regimen. Exercise specifically elevated copper levels in the GA muscle compared to sedentary controls (Figure 1A), whereas other transition metals–including iron, zinc, and manganese–remained unchanged (Figure S1). To determine whether this targeted copper accumulation is functionally required for muscle performance, we disrupted local copper acquisition by generating skeletal muscle-specific knockout of the primary cellular copper transporter, *Ctr1* (SMKO), by crossing *Ctr1*-floxed mice (Floxed) with myosin light chain 1 (*Myl1*)-Cre transgenic mice. Following validation of *Ctr1* genomic excision and transcript depletion (Figures S2A-S2C), we confirmed that SMKO muscles exhibited significantly diminished basal copper pools (Figure 1B). Although SMKO mice maintained body weight and skeletal muscle mass comparable to Floxed controls (Figures S2D and S2E), they displayed marked deficits in forelimb grip strength (Figure 1C) and endurance exercise capacity (Figure 1D). These results establish that CTR1-mediated copper uptake is essential for maintaining skeletal muscle strength and exercise tolerance.

**Figure 1.**
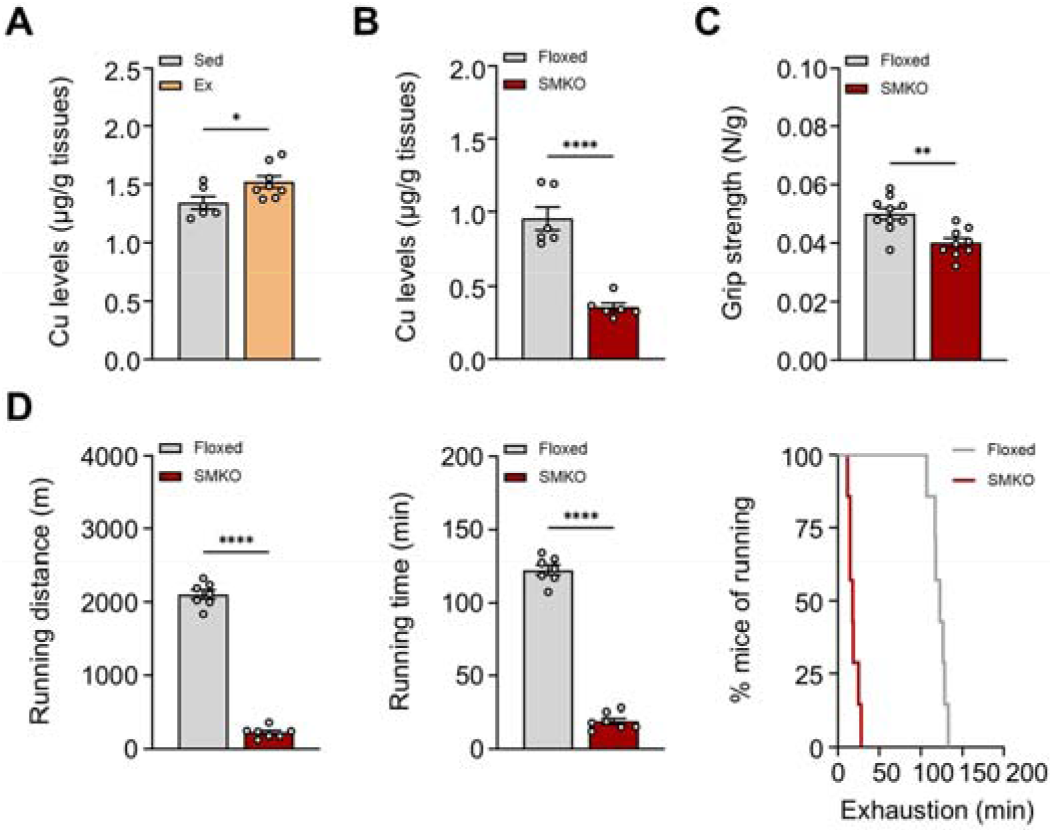
Ctr1-mediated copper uptake supports skeletal muscle performance. (A and B) Copper levels in the GA skeletal muscle of exercise-trained wild-type mice (A) and SMKO mice (B) compared to respective controls (*n* = 6–8). (C) Forelimb grip strength of Floxed and SMKO mice (*n* = 9–10). (D) Running distance, running time, and exhaustion rate of Floxed and SMKO mice during treadmill training (*n* = 7). Data are presented as mean ± SEM. Statistical differences were determined by two-tailed Student’s t-test.

### Skeletal muscle copper deficiency restricts systemic and cellular oxygen consumption, enforcing a reliance on glycolysis

Given the endurance deficit in SMKO mice, we examined skeletal muscle fiber type composition, as slow-twitch (type I) fibers typically confer fatigue resistance.^14^ Paradoxically, SMKO muscles exhibited macroscopic reddening (Figure 2A) and a robust shift toward type I and type IIa fibers, as confirmed by myosin heavy chain (MyHC) immunostaining (Figure 2B). This transition was corroborated at the transcriptional level by the upregulation of *Myh7* (type I) and *Myh2* (type IIa), concomitant with the downregulation of fast-twitch markers *Myh1* (type IIx) and *Myh4* (type IIb) (Figure 2C). Consistent with this slow-twitch enrichment, SMKO myofibers displayed a reduced cross-sectional area (CSA) (Figure 2D). Because histological assessment revealed no overt atrophy or tissue degeneration (Figure S3A), we conclude that this reduced CSA reflects the natural morphological transition to smaller-caliber slow-twitch fibers rather than pathological wasting.^25^

**Figure 2.**
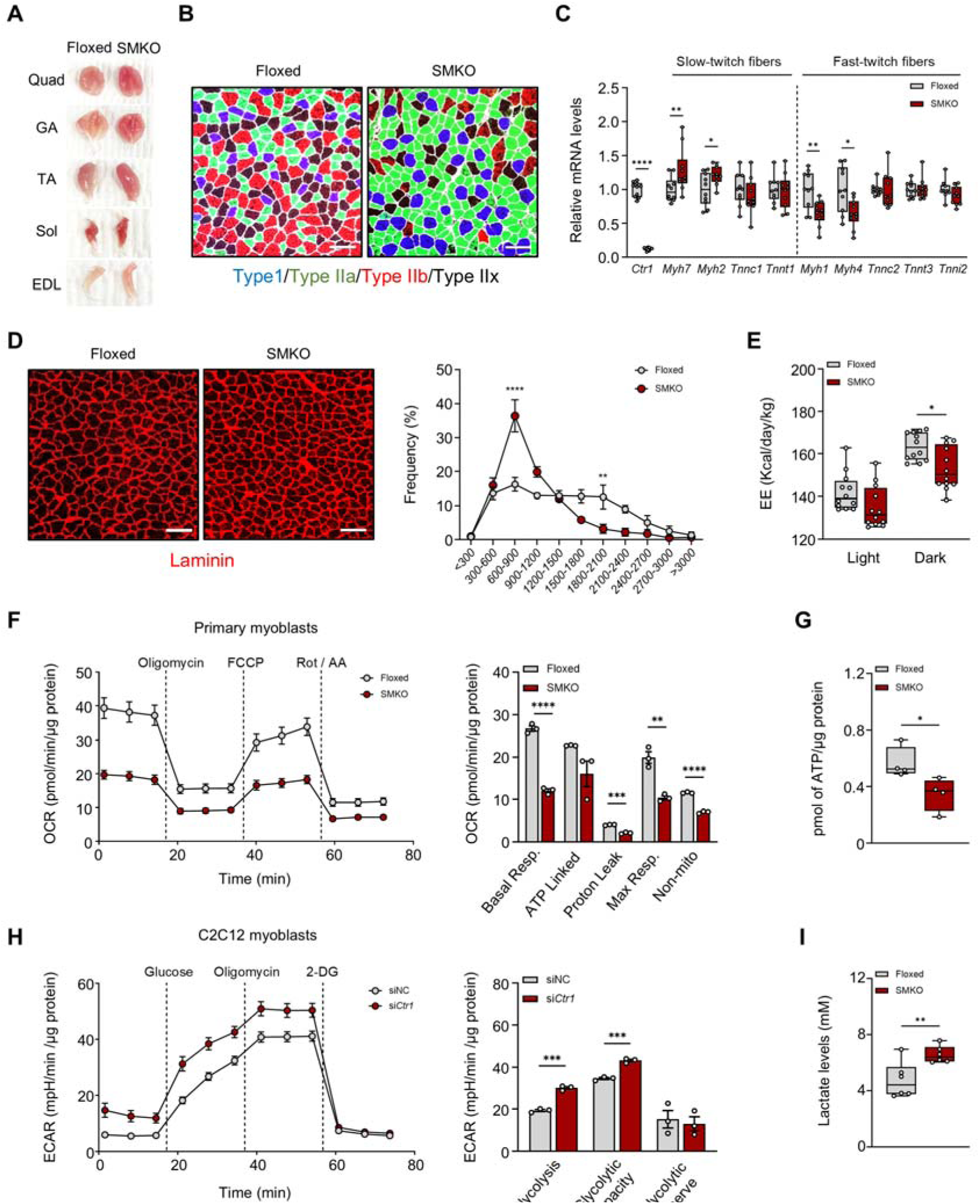
Skeletal muscle copper deficiency suppresses systemic energy expenditure and cellular oxygen consumption, forcing a glycolytic shift. (A) Representative gross macroscopic images of skeletal muscle from Floxed and SMKO mice. (B) Representative immunofluorescence staining of myofiber types in the GA muscle. Scale bars = 100 μm. (C) Relative mRNA expression of slow- and fast-twitch fiber-related genes in the GA muscle (*n* = 10). (D) Cross-sectional area (CSA) of each myofiber in the GA muscle of Floxed and SMKO mice. CSA frequency was calculated using ImageJ. Scale bars = 100 μm. (E) Total energy expenditure evaluated via indirect calorimetry. (F) Oxygen consumption rate (OCR) in primary myoblasts isolated from the hindlimb muscles of Floxed and SMKO mice, measured following sequential addition of oligomycin (Oligo), carbonyl cyanide p-triflouromethoxyphenylhydrazone (FCCP), rotenone (Rot), and antimycin A (AA). (G) Steady-state ATP levels in the GA muscle (*n* = 4). (H) Extracellular acidification rate (ECAR) in C2C12 myoblasts following siRNA-mediated knockdown (*NCi*, negative control siRNA; *Ctr1i*, *Ctr1* siRNA). (I) Circulating blood lactate concentrations in Floxed and SMKO mice (*n* = 6). Data are presented as mean ± SEM. Statistical differences were determined by two-way ANOVA with Šídák’s multiple comparisons test (E) or two-tailed Student’s t-test (C and F-I).

Because slow-twitch fiber enrichment typically elevates systemic energy expenditure and endurance,^14^ we evaluated whole-body metabolism using indirect calorimetry. Despite the structural shift toward oxidative fiber types, SMKO mice exhibited significantly reduced oxygen consumption (VO□) (Figure S3B) without corresponding changes in carbon dioxide production (VCO□) (Figure S3C), culminating in suppressed total energy expenditure (Figure 2E). To determine whether this systemic phenotype stemmed from intrinsic bioenergetic defect,^26^ we measured respiratory flux in primary myoblasts isolated from SMKO mice. *Ctr1* depletion induced a global suppression of mitochondrial respiration, with significant reductions observed across all respiratory states (Figure 2F). This broad attenuation of respiratory capacity directly translated to the *in vivo* setting, where steady-state ATP levels were significantly diminished in SMKO GA muscle (Figure 2G). To investigate whether this mitochondrial failure triggers metabolic reprogramming, we quantified the extracellular acidification rate (ECAR) in *Ctr1*-knockdown C2C12 myoblasts. Copper depletion significantly elevated glycolytic flux (Figure 2H), a metabolic shift that was mirrored *in vivo* by elevated circulating lactate concentrations in SMKO mice (Figure 2I). Together, these findings demonstrate that skeletal muscle copper deficiency severely compromises mitochondrial respiration, rendering the structural slow-twitch fiber transition functionally ineffective and forcing a compensatory, yet insufficient, reliance on glycolysis.

### Skeletal muscle copper deficiency alters ETC subunit composition and induces mitochondrial myopathy phenotype

To define the molecular basis of the bioenergetic collapse observed in SMKO muscle, we performed a proteomic analysis of GA tissue from Floxed and SMKO mice. Using a fold-change cutoff of 1.5 and an adjusted *p* value < 0.05, we identified 88 differentially expressed proteins (DEPs), comprising 53 upregulated and 35 downregulated proteins in SMKO muscle (Figure 3A). A substantial proportion of DEPs were mitochondrial proteins directly associated with OXPHOS (Figures 3B and 3C). Gene ontology (GO) analysis revealed that the downregulated proteome in SMKO muscle was heavily enriched for components of ETC Complex IV, particularly proteins mediating electron transfer from cytochrome c to oxygen (Figure 3D). Strikingly, the most significantly downregulated proteins were cytochrome c oxidase subunit 1 (MTCO1/COX1) and MTCO2/COX2–critical copper-binding subunits essential for Complex IV assembly and function (Figure 3E).^27^ In contrast, upregulated proteins were highly enriched for pathways related to Complexes I and II, as well as ATP synthesis, suggesting a compensatory response to the severe Complex IV depletion (Figures 3E and S4). Western blot analysis corroborated this asymmetric ETC imbalance–revealing a profound depletion in Complex IV alongside the compensatory upregulation of other ETC complexes–while the concomitant induction of the intracellular sensor copper chaperone for superoxide dismutase (CCS) confirmed the underlying functional copper depletion (Figure 3F).^28,29^

**Figure 3.**
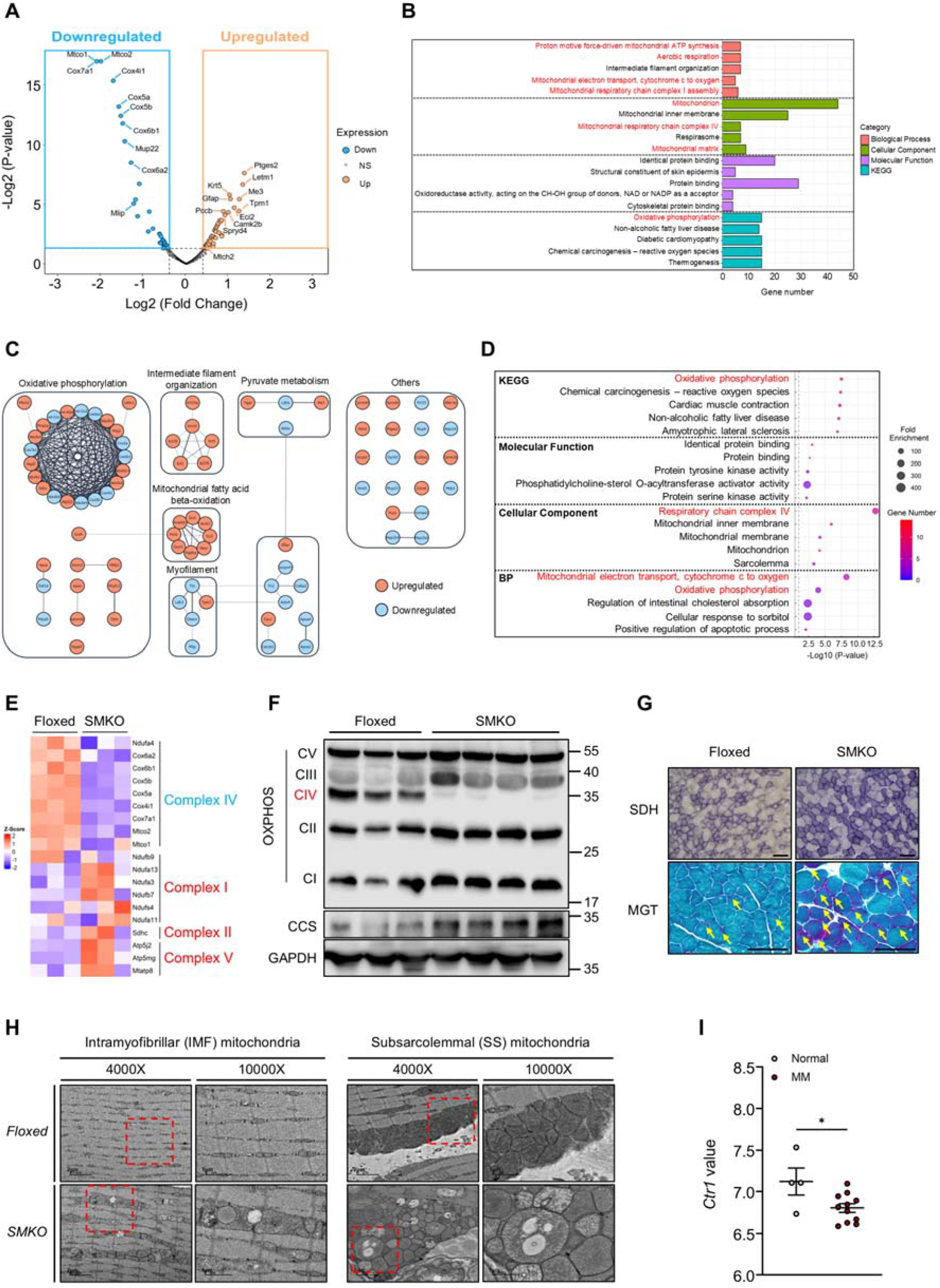
Copper deficiency induces Complex IV depletion, compensatory ETC remodeling, and hallmarks of MM. (A-D) Proteomics analysis of GA muscle from Floxed and SMKO mice. (A) Volcano plot. (B) GO analysis of DEPs. (C) STRING network analysis of DEPs. (D) GO analysis of downregulated proteins in SMKO compared to Floxed mice. (E) Heatmap visualization of OXPHOS-related protein expression. (F-H) Analysis of mitochondrial function in the GA muscle of Floxed and SMKO mice. (F) Protein levels of OXPHOS components and CCS. (G) SDH and modified Gomori trichorme (MGT) staining. Scale bar = 100 μm. (H) Transmission electron microscopy (TEM) images. Scale bar = 2 μm (4,000× magnification), 1 μm (10,000× magnification). (I) *CTR1* expression levels of individuals with the MM compared to normal controls (extracted from GSE43698). Data are presented as mean ± SEM. Statistical differences were determined by two-tailed Student’s t-test (I).

Mitochondrial myopathy (MM) is characterized by mitochondrial dysfunction and is frequently associated with a highly specific molecular signature: Complex IV deficiency coupled with compensatory Complex II upregulation and elevated circulating lactate.^16,30, 31^ Given the striking phenotypic and proteomic parallels, we hypothesized that *Ctr1* deletion in skeletal muscle provokes a pathology mimicking clinical MM. Histological and ultrastructural evaluations strongly supported this; SMKO mice exhibited severe MM-like features, including robust succinate dehydrogenase (SDH) hyperactivation, the widespread appearance of ragged-red fibers, and enlarged, morphologically aberrant mitochondria (Figures 3G and 3H). To determine the clinical relevance of this molecular pathology, we analyzed a human GEO dataset (GSE43698) and found that *CTR1* transcript levels were significantly reduced in muscle biopsies from MM patients compared to healthy controls (Figure 3I).

To examine whether the observed metabolic and histopathological changes are sex-dependent, we evaluated exercise performance and muscle pathology in female SMKO mice. Consistent with the male cohort, female SMKO mice exhibited severely restricted exercise capacity (Figures S5A and S5B) alongside elevated SDH activity and the presence of ragged-red fibers. Interestingly, female SMKO did not exhibit elevated basal blood lactate levels (Figures S5C and S5D), suggesting that while the overarching histological and functional hallmarks of MM are sex-independent, the degree of systemic glycolytic spillover may be influenced by sex-specific metabolic buffering. Together, these findings demonstrate that copper deficiency drives a profound compositional imbalance within the ETC, inducing both the molecular and histological hallmarks of MM.

### Copper directly binds to MTCH2 to govern its proteasomal turnover

Building on our proteomic analysis of SMKO muscle, we sought to identify candidate copper-binding proteins that might drive mitochondrial dysfunction under copper starvation. Among the 88 DEPs, interrogation utilizing the metal-binding prediction tool MeBiPred^32^ identified six potential copper-binding candidates, including the canonical COX subunits MTCO1 and MTCO2 (Figure S6A). We focused our investigation on mitochondrial carrier homolog 2 (MTCH2, SLC25A50) due to its predicted Cu^+^ specificity, OMM localization, and previously defined roles in mitochondrial dynamics, apoptosis, and OXPHOS reguation.^33–35^ *In vitro* metal-binding assays revealed that recombinant MTCH2 selectively bound to copper-charged resin with dose-dependent affinity, while showing no interaction with iron-, zinc-, or uncharged resins, (Figure 4A). To investigate the structural basis of this interaction, we modeled the theoretical conformation of MTCH2 using AlphaFold 3, as its experimental structure remains unresolved. The predicted model features an open α-barrel alongside a bulky region comprising several α-helices and two β-strands located at one end of the α-barrel (Figure 4B). Polar and hydrophilic residues line the barrel’s interior, while numerous hydrophobic residues are externally exposed, consistent with its integration in the lipid bilayer.^36^ Notably, this model revealed that the barrel’s internal surface is unusually exposed to the lipid phase (Figure 4B, left panel, red dotted circle)–a structural feature rarely observed in experimentally determined membrane proteins. To identify a closed α-barrel with analogous folding, we conducted a structural similarity search using the DALI server. Using the AlphaFold-predicted MTCH2 as a query retrieved highly homologous structures (PDB ID 8J1N, mitochondrial brown fat uncoupling protein; PDB IDs 1OKC and 4C9H, mitochondrial ADP/ATP carrier proteins) with Z-scores exceeding 20.^37,38^ These homologs superimpose on the AlphaFold-predicted open MTCH2 structure with a root-mean-square deviation (RMSD) of less than 3.1 Å, featuring a closed α-barrel surrounded by flexible regions and associated lipids, characteristic of integral membrane transporters. Using the ADP/ATP carrier protein (PDB ID 1OKC, SLC25A4) as a template, a refined MTCH2 structure was generated via SWISS MODEL.^39^ In this closed conformation, the open α-barrel is sealed by the incorporation of a C-terminal helix (N282-Y295, designated the C-helix) (Figure 4B, middle panel). By integrating primary sequence features with this refined structural modeling, we identified two discrete candidate copper-binding interfaces: an exterior cluster of sulfur-rich residues (M244, C268, M278) and an interior lining of nitrogen- and oxygen-rich residues (H142, R185, S241; H88, E127, R131) (Figure 4B, right panel). To functionally validate these predicted sites, we generated alanine substitution mutants and evaluated their copper-binding capacity. Strikingly, the M244A, C268A, and M278A mutants exhibited profoundly diminished copper binding, establishing these exterior sulfur-rich residues as critical mediators of the MTCH2-copper interaction (Figures 4C and S6B).

**Figure 4.**
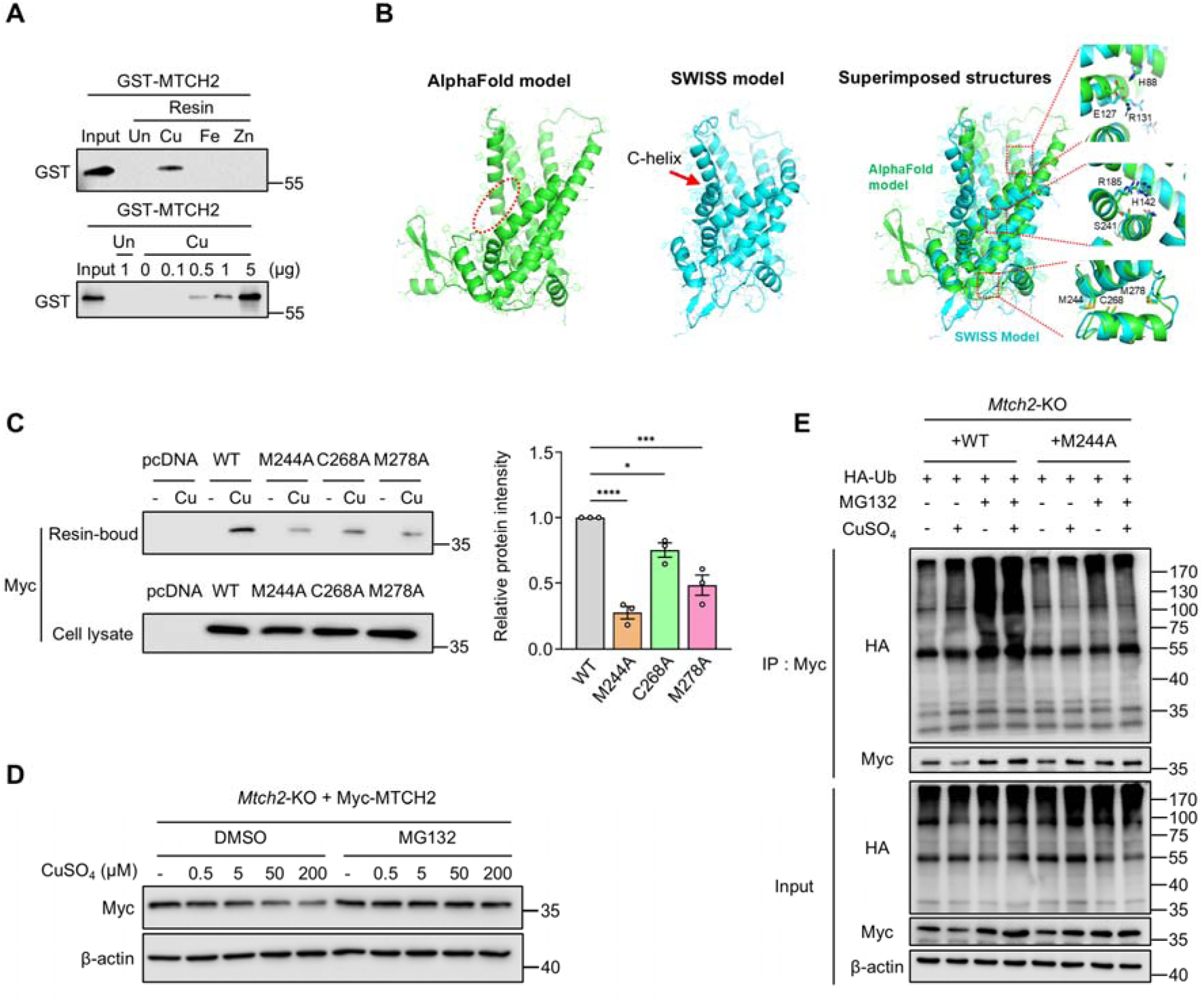
MTCH2 stability is dictated by direct copper binding and proteasomal clearance. (A) *In vitro* copper-binding assay utilizing glutathione S-transferase (GST)-tagged human recombinant MTCH2 protein against transition metal-charged resins. (B) Predicted structural models of MTCH2 generated by Alphafold3 (left panel) and SWISS-MODEL (middle panel). Candidate amino acids clusters mediating copper binding are detailed (right panel). (C) *In vitro* copper-binding validation in HEK293T cells expressing wild-type (WT) or alanine-substitution mutants pCMV3-MTCH2-Myc. Bound protein fractions were normalized to total Myc expression in whole-cell lysates. *n* = 3 independent experiments. (D) Immunoblot analysis detailing MTCH2 stability in *Mtch2*-knockout C2C12 myoblasts re-expressing MTCH2-WT or the copper-binding deficient MTCH2-M244A. Cells were pre-treated with the proteasome inhibitor MG132 (1μM) for 2 h prior to CuSO_4_ exposure (200 μM, 6 h). (E) Co-immunoprecipitation assay assessing MTCH2 polyubiquitination. *Mtch2*-knockout C2C12 myoblasts co-transfected with HA-ubiquitin and MTCH2-WT or MTCH2-M244A were pre-treated with MG132 (1μM, 2 h) followed by CuSO_4_ treatment (250 μM, 6 h). Data are presented as mean ± SEM. Statistical differences were determined by one-way ANOVA with Tukey’s test (C).

Given that intracellular copper availability frequently dictates protein half-life, and MTCH2 was ∼1.8-fold upregulated in SMKO muscle (Figure S6A), we next examined whether MTCH2 undergoes copper-dependent turnover. In *Mtch2-*KO C2C12 myoblasts re-expressing wild-type MTCH2 (MTCH2-WT), CuSO_4_ treatment induced a dose-dependent reduction in MTCH2 stability (Figure 4D). This degradation was effectively rescued by MG132, indicating proteasome-mediated clearance mechanism.^40^ Importantly, excess copper stimulated the polyubiquitination of MTCH2-WT, a modification that further accumulated upon proteasomal inhibition. In contrast, copper-binding deficient M244A mutant failed to undergo ubiquitination and remained entirely resistant to copper-induced degradation (Figure 4E). Collectively, these results demonstrate that MTCH2 is a direct copper-binding protein, and that copper coordination at specific sulfur-rich residues acts as a molecular switch governing its polyubiquitination and proteasomal clearance.

### MTCH2 integrates mitochondrial dynamics with copper trafficking

Given that copper directly dictates MTCH2 stability, we next examined whether its functional roles are governed by copper availability. Given MTCH2’s established role in driving mitochondrial fusion,^33,41^ the structural hyperfusion observed in SMKO muscle suggested a mechanistic link between copper starvation and MTCH2-medated fusion. Indeed, mitochondria isolated from SMKO quadriceps (Quad) exhibited pronounced accumulation of MTCH2 alongside the canonical fusion proteins MFN2 and OPA1, with no compensatory alterations in fission markers (Figure 5A). To confirm that MTCH2 actively drives this structural remodeling, we mimicked copper deficiency *in vitro* using the specific copper chelator bathocuproinedisulfonic acid (BCS), which induced widespread mitochondrial elongation (Figure S7A).^20^ As expected, *Mtch2* silencing in C2C12 myoblasts was sufficient to induce baseline mitochondrial fragmentation.^33,41^ Crucially, this depletion completely abrogated BCS-induced elongation (Figure 5B), demonstrating that MTCH2 is strictly required for the structural hyperfusion triggered by copper starvation. Conversely, we assessed whether copper overload–which accelerates MTCH2 degradation–would compromise mitochondrial fusion. In *Mtch2*-KO C2C12 myoblasts, re-expression of either MTCH2-WT or the copper-binding-deficient M244A mutant effectively rescued mitochondrial elongation, indicating that direct copper binding is dispensable for MTCH2’s core fusion activity (Figure 5C). However, under copper overload, MTCH2-WT underwent rapid degradation, resulting in the collapse of the network into fragmented mitochondria. In striking contrast, the M244A mutant resisted degradation and preserved an elongated mitochondrial architecture (Figure 5C). These findings establish that copper coordination at M244 serves as a critical structural switch determining MTCH2 half-life, thereby exerting secondary control over mitochondrial morphology.

**Figure 5.**
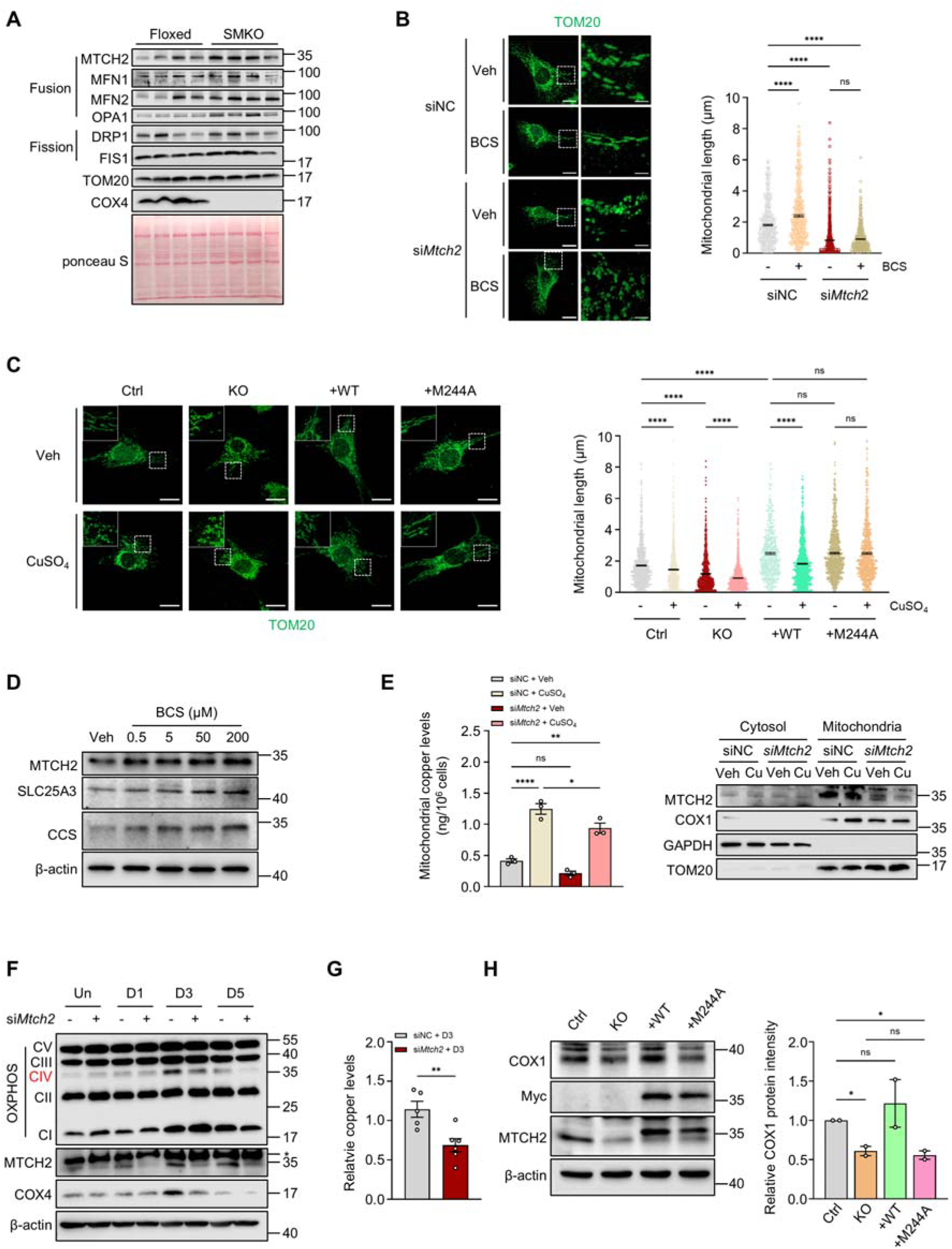
MTCH2 integrates mitochondrial dynamics with copper trafficking. (A) Immunoblot analysis of mitochondria isolated from the Quad muscle of Floxed and SMKO mice. (B and C) Immunofluorescence imaging of mitochondrial morphology in *Mtch2* knockdown C2C12 myoblasts treated with the copper chelator BCS (50 μM) (B), and *Mtch2* knockout C2C12 myoblasts re-expressing MTCH2-WT or MTCH2-M244A following CuSO_4_ exposure (200 μM, 6 h) (C). Scale bars = 20 μm (main) and 5 μm (inset). Mitochondrial length was quantified from ∼1,000 mitochondria across 10-15 cells per condition using ImageJ. (D) Immunoblot analysis of human skeletal muscle myoblasts (HSMM) treated with vehicle or BCS (50 μM, 24 h). (E) ICP-MS quantification of mitochondrial copper (left) and parallel immunoblot analysis of cytosolic and mitochondrial fractions (right) in *Mtch2*-knockdown C2C12 myoblasts exposed to CuSO_4_ (50 μM) for 24 h. siNC, negative control siRNA; si*Mtch2*, *Mtch2* siRNA. *n* = 3. (F) Immunoblot analysis of differentiated C2C12 myotubes following targeted *Mtch2* knockdown. (G) ICP-MS analysis of mitochondrial copper accumulation over 3 days of myogenic differentiation in C2C12 cells following *Mtch2* knockdown. (H) Immunoblot analysis of differentiated *Mtch2*-knockout C2C12 myotubes re-expressing MTCH2-WT or MTCH2-M244A, treated with CuSO_4_ (50 μM) for 24 h. COX1 protein expression was normalized to β-actin. *n* = 2 independent experiments. Data are presented as mean ± SEM. Statistical differences were determined by two-tailed Student’s t-test (G) or one-way ANOVA (B, C, E and H) with Tukey’s test.

Beyond its role in membrane dynamics, MTCH2 localizes to the OMM and belongs to the SLC25 family of metabolite carriers.^42,43^ Because intracellular metal overload frequently triggers the targeted degradation of transporters to prevent cellular toxicity,^44^ we hypothesized that MTCH2 might directly facilitate mitochondrial copper uptake. Supporting this, both MTCH2 and the inner-membrane copper transporter SLC25A3 were upregulated by BCS-induced copper depletion across multiple cell lines (Figures 5D and S7B). Furthermore, silencing *Mtch2* significantly restricted mitochondrial copper pools under copper-replete–but not basal–conditions (Figure 5E), and severely impaired both Complex IV expression and mitochondrial copper accumulation during myogenic differentiation (Figure 5F and 5G). Strikingly, while re-expression of MTCH2-WT in *Mtch2*-KO myotubes successfully restored COX1 protein levels, the copper-binding deficient M244A mutant failed to rescue COX1 expression (Figure 5H). Together, these findings genetically dissociate MTCH2’s dual function: its physical accumulation drives adaptive mitochondrial hyperfusion during copper starvation, while its distinct copper-binding capacity is required to traffic copper across the OMM to sustain Complex IV activity. Ultimately, these integrated roles identify MTCH2 as a central metabolic sensor that couples cellular copper status to both mitochondrial architecture and respiratory competence.

### The copper ionophore elesclomol restores mitochondrial function and exercise capacity in SMKO mice

Elesclomol (ES) is a lipophilic copper ionophore that facilitates intracellular copper delivery independently of CTR1. By delivering bioavailable copper directly to intracellular compartments, ES has been shown to support COX activity in copper-deficient cellular models and reduce mortality in a mouse model of Menkes disease.^47^ These findings raised the possibility that targeted, pharmacological copper supplementation could bypass the sarcolemmal uptake defect in SMKO mice and reverse the resulting MM.

To test this, we administered ES subcutaneously to Floxed and SMKO mice for 2-3 weeks. ES treatment restored intracellular copper availability, significantly increasing copper levels in the GA muscle (Figure 6A). This biochemical rescue translated to a profound physiological recovery: ES substantially improved exercise capacity (distance and duration) in SMKO mice (Figure 6B) and reversed the pathological shift toward slow-twitch fibers. This normalization of muscle phenotype was evident from both macroscopic tissue color and immunostaining of myofiber composition (Figures S8A and 6C). Critically, this histological rescue was accompanied by the reversal of the bioenergetic collapse observed in the knockout model. ES treatment enhanced mitochondrial respiration in primary myoblasts isolated from SMKO mice and significantly reduced blood lactate concentrations (Figures 6D and 6E), indicating a successful reversal of the maladaptive glycolytic reprogramming. In line with these findings, ES normalized whole-body VO_2_ and total energy expenditure (Figures S8B and 6F).

**Figure 6.**
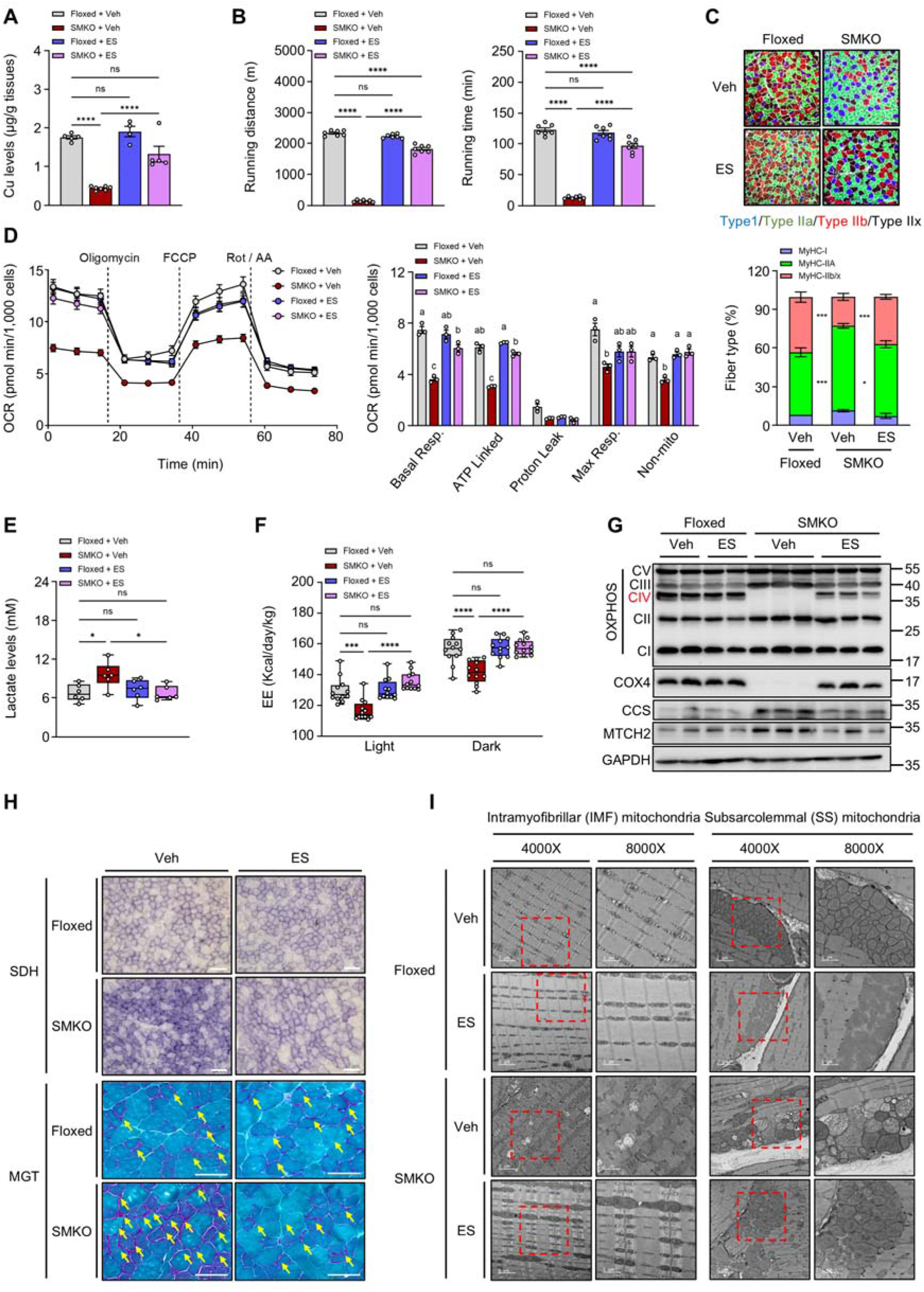
The copper ionophore elesclomol restores mitochondrial function and exercise capacity in SMKO mice. ES was administrated subcutaneously to Floxed and SMKO mice for 2-3 weeks. (A) Total copper levels in the GA muscle (*n* = 4–7). (B) Exercise capacity measured by treadmill running distance and duration (*n* = 7). (C) Immunofluorescence staining and quantification of myofiber type composition. (D) OCR of isolated primary myoblasts. Bars labeled with distint letters (a, b, c, d) indicate statistically significant differences between groups (*p* < 0.05). (E) Blood lactate concentration (*n* = 6). (F) Whole-body energy expenditure (*n* = 5–7). (G) Representative immunoblots of OXPHOS complexes, CCS, COX4, and MTCH2 protein levels. (H) Histopathological evaluation via SDH and MGT staining. Scale bar = 100 μm. (I) Representative TEM images. Scale bar = 2 μm (4,000× magnification), 1 μm (8,000× magnification). Data are presented as mean ± SEM. Statistical differences were determined by one-way ANOVA (A, B, D and E) or two-way ANOVA (C and F) with Tukey’s test.

At the molecular level, ES treatment restored the compositional balance of the ETC by rescuing Complex IV expression. Furthermore, ES normalized the expression of MTCH2 and CCS, confirming that intracellular copper sensing was reset to baseline (Figure 6G). Finally, ES administration ameliorated severe histopathological features, resolving the appearance of ragged-red fibers (SDH and MGT staining) and rescuing mitochondrial ultrastructural abnormalities (Figures 6H and 6I). Together, these findings demonstrate that pharmacological copper delivery via ES can fully bypass the genetic deletion of *Ctr1*, successfully reversing the ETC imbalance and MM-like phenotypes in SMKO mice.

### AAV-DIO-mediated *Ctr1* restoration reverses mitochondrial myopathy and metabolic reprogramming in SMKO mice

To definitively establish that the MM-like pathology in SMKO mice is driven by sarcolemmal copper transport deficiency, we utilized an AAV9-based gene delivery approach. We employed a Cre-activated AAV-double-floxed inverted open reading frame (DIO) system to selectively reinstate *Ctr1* expression in skeletal muscle (Figure S9A). Following *in vivo* AAV-DIO-*Ctr1* administration, *Ctr1* expression was successfully restored in SMKO muscle (Figure S9B), which effectively rescued intracellular copper pools (Figure 7A). This genetic restoration translated into profound physiological improvements, including normalized treadmill running capacity (Figure 7B). Furthermore, targeted Ctr1 replacement reversed the pathological shift toward slow-twitch fibers, returning the muscle phenotype to baseline as evidenced by both macroscopic tissue color and myofiber immunostaining profiles (Figures S9C and 7C). The reestablishment of copper uptake also resolved the systemic and cellular bioenergetic collapse: it enhanced mitochondrial respiration in isolated primary myoblasts (Figure 7D), decreased blood lactate accumulation (Figure 7E), and normalized whole-body VO_2_ and total energy expenditure (Figures S9D and 7F). Crucially, restoring the intracellular copper pool reset the MTCH2-mediated starvation response. MTCH2 and CCS protein levels were normalized, accompanied by the rescue of Complex IV expression (Figure 7G). Histological and ultrastructural analyses further confirmed the clearance of ragged-red fibers and the restoration of normal mitochondrial architecture (Figures 7H and 7I). Collectively, these findings demonstrate that targeted *Ctr1* re-expression is sufficient to re-establish skeletal muscle copper homeostasis and fully reverse the metabolic and mitochondrial defects of copper-deficiency myopathy.

**Figure 7.**
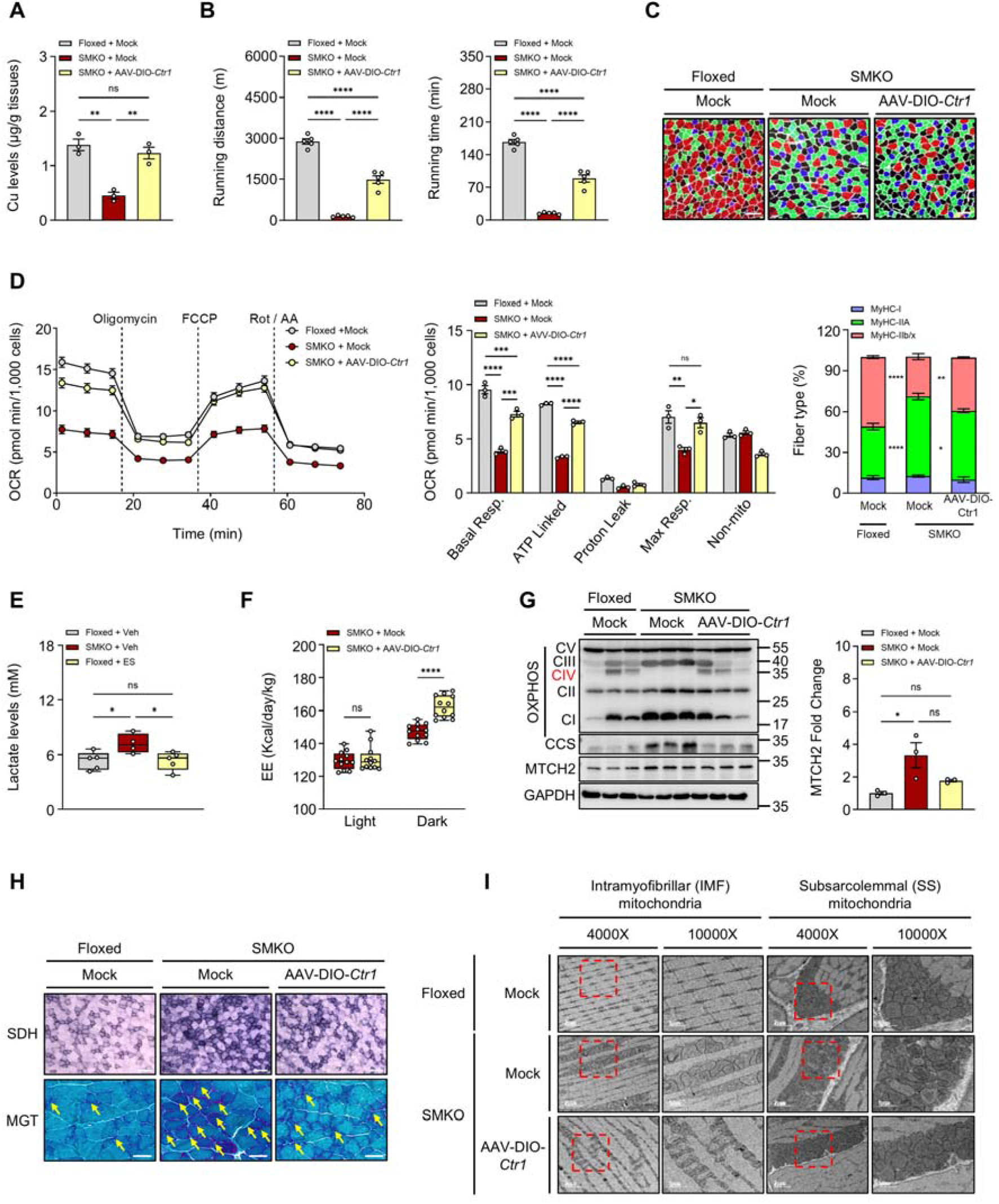
AAV-DIO-mediated *Ctr1* re-expression in skeletal muscle restores mitochondrial function and improves exercise performance in SMKO mice. A single intramuscular injection of AAV-DIO-*Ctr1* was administered to SMKO mice, and assessments were performed after 5 weeks. (A) Copper levels in the GA muscle (*n* = 3). (B) Running distance and duration during treadmill exercise (*n* = 5). (C) Immunofluorescence staining of myofiber types. (D) OCR of primary myoblasts. (E) Blood lactate concentration (*n* = 4–5). (F) Whole-body energy expenditure. (G) OXPHOS, CCS, and MTCH2 protein levels, normalized to GAPDH expression. (H) SDH and MGT staining. Scale bar = 100 μm (SDH) or 50 μm (MGT). (I) Representative TEM images of mitochondrial ultrastructure. Scale bar = 2 μm (4,000× magnification), 1 μm (10,000× magnification). Data are presented as mean ± SEM. Statistical differences were determined by one-way ANOVA (A, B, E and G) and two-way ANOVA with Tukey’s test (C and D) or Šídák’s multiple comparisons test (F). ns, not significant.

## DISCUSSION

Mitochondrial function is fundamental to skeletal muscle metabolism, and defects in OXPHOS contribute substantially to muscle weakness and systemic metabolic disorders.^48^ Copper, an essential cofactor for COX in Complex IV of the ETC, is indispensable for mitochondrial respiration, yet its regulatory pathway and precise physiological roles in adult skeletal muscle have remained poorly understood. Here, we identify a critical role for intracellular copper availability in maintaining mitochondrial integrity and whole-body muscle performance, mediated in part by the OMM protein MTCH2. Skeletal muscle–specific deletion of the primary copper importer CTR1 provoked progressive copper starvation, culminating in a MM-like phenotype characterized by severe exercise intolerance, ETC proteome asymmetry, bioenergetic collapse, and systemic metabolic reprogramming.

Mitochondrial copper homeostasis has been primarily attributed to the inner membrane carrier SLC25A3, which imports copper into the matrix to sustain COX activity.^49,50^ While it was postulated that copper traverse the OMM when bound to apoligand,^51^ the molecular mediators of this process remained elusive. Our analysis identified MTCH2–an OMM resident SLC25 family–as a highly responsive, differentially regulated copper-binding protein in SMKO mice. We demonstrate that MTCH2 directly coordinates copper via a motif containing a sulfur-rich amino acid cluster centered on the M244 residue. Crucially, MTCH2 deletion depleted mitochondrial copper content and diminished COX expression in myoblasts, while copper-binding-deficient MTCH2 mutants (M244A) failed to restore COX1 levels. These data firmly establish MTCH2 as a critical mediator of mitochondrial copper trafficking.

Beyond canonical trafficking, our data reveal that intracellular copper status actively dictates mitochondrial architecture, coopting MTCH2’s previously established role in mitochondrial dynamics. Under basal conditions, adequate copper sustains COX activity and promotes steady-state mitochondrial membrane potential “flickering” –transient depolarization events that activate the metalloendopeptidase OMA1, driving OPA1 cleavage and thereby restraining excessive fusion.^52^ Under copper-deficient conditions, however, this flickering is suppressed and OPA1 remains intact. Simultaneously, copper starvation stabilizes the MTCH2 protein. Consequently, mitochondria undergo massive hyperfusion. Importantly, this ultrastructural shift is completely abolished by MTCH2 knockdown, demonstrating that MTCH2 is strictly required to drive hyperfusion under copper deficiency. Conversely, copper overload destabilizes MTCH2, reducing its pro-fusion activity and inducing mitochondrial fragmentation. Together, these observations unveil a previously unrecognized dual function for MTCH2, acting simultaneously as a copper trafficker and a copper-sensitive regulator of mitochondrial fusion.

This dual functionality elegantly resolves a prevailing paradox in the literature. Previous work reported that skeletal muscle-specific MTCH2 deletion enhances mitochondrial respiration, promotes oxidative fiber-type switching, and improves exercise performance.^35,53^ At first glance, this appears to contradict our finding that MTCH2 facilitates copper delivery to Complex IV. However, we propose that these divergent phenotypes reflect the presence of basal compensatory mechanisms and the context-dependent nature of MTCH2. Under steady-state conditions, alternative pathways (such as robust CTR1-mediated uptake) likely preserve sufficient mitochondrial copper pools even in the absence of MTCH2, thereby sustaining COX activity and flickering. Consequently, the isolated loss of MTCH2’s pro-fusion function dominates the phenotype, resulting in mitochondrial fragmentation and subsequent metabolic compensations. By contrast, in our SMKO model, the upstream deletion of CTR1 universally restricts cellular copper availability. Under these severe starvation conditions, the intracellular copper pool is exhausted, OXPHOS collapses, and MTCH2 is pathologically stabilized to drive unchecked hyperfusion. These contrasting phenotypes highlight that MTCH2’s contribution to copper homeostasis becomes biologically critical primarily when cellular copper flux is restricted. Future investigations incorporating direct quantification of mitochondrial copper pools and isolated COX activity in MTCH2 KO muscle will be essential to fully disentangle these dual influences.

Our findings also underscore the broader physiological necessity of dynamic copper regulation in mature skeletal muscle. While copper has previously been implicated in myoblast proliferation and differentiation,^23,24^ our study extends this paradigm, demonstrating that active copper compartmentalization is essential for sustaining metabolic and structural integrity in adult tissue. This has profound implications for aging and muscle-related pathologies, including MM and sarcopenia, where metabolic decline and progressive ETC dysfunction are common features.^31,54^ Furthermore, reduced exercise capacity is a hallmark of Menkes disease–a systemic copper deficiency disorder caused by ATP7A mutations^47^–and is observed in amyotrophic lateral sclerosis models harboring superoxide dismutase [Cu-Zn] mutations.^55,56^ In these diverse disease contexts, disrupted copper compartmentalization likely represents a fundamental driver of endurance impairment and mitochondrial failure.

Therapeutically, our data demonstrate that restoring copper availability effectively reverses mitochondrial and metabolic dysfunction in copper-deficient muscle. Treatment with ES, a copper ionophore that bypasses CTR1-mediated transport, successfully restored intracellular copper levels, rescued OXPHOS activity, and improved exercise capacity in SMKO mice. Likewise, AAV-mediated re-expression of *Ctr1* restored metabolic balance and mitochondrial function. With AAV9 therapies already in clinical use for neuromuscular disorders and expanding interest in targeting metabolic myopathies,^16,57–59^ manipulating the copper-mitochondria axis represents a highly promising, underexplored therapeutic avenue.

In conclusion, this study identifies CTR1 and MTCH2 as essential regulators of mitochondrial copper homeostasis and skeletal muscle function. We reveal MTCH2 as a critical dual-function protein that not only coordinates copper delivery to ETC but also integrates cellular copper status with mitochondrial dynamics. These findings establish a previously unrecognized mechanistic link between copper metabolism, mitochondrial architecture, and muscle performance, offering new translational perspectives for the treatment of muscular disorders driven by copper dyshomeostasis.

### Limitations of the study

Our study underscores the crucial roles of CTR1 and MTCH2 in maintaining mitochondrial copper homeostasis and skeletal muscle function. We identified MTCH2 as a copper-binding mitochondrial protein and proposed that it regulates mitochondrial dynamics and respiratory function in response to cellular copper status. However, it remains unclear whether MTCH2 directly transports cytosolic copper into mitochondria across the OMM or instead indirectly modulates mitochondrial copper availability. Additional studies are also needed to determine how cytosolic copper is delivered to MTCH2. Elucidating this distinction will require copper flux assays and high-resolution structural analysis of copper-bound MTCH2. Furthermore, how MTCH2 interacts with mitochondrial inner membrane copper transporters such as SLC25A3 or with copper chaperones such as COX17, and whether these functions are conserved across tissues and species, remains to be investigated. Lastly, while ES treatment and AAV-mediated *Ctr1* restoration showed therapeutic potential, future studies are needed to explore whether modulating copper homeostasis could broadly benefit other muscle disorders involving mitochondrial dysfunction.

## RESOURCE AVAILABILITY

### Lead contact

Further information and requests for resources and reagents should be directed to and will be fulfilled by the lead contact, Tae-Il Jeon.

### Materials availability

All unique reagents generated in this study are available from the lead contact with a completed material transfer agreement.

### Data and code availability

The mass spectrometry proteomics data have been deposited to the ProteomeXchange Consortium via the PRIDE partner repository with the dataset identifier PXD064414

## ACKNOWLEDGMENTS

This work was supported by the National Research Foundation of Korea (NRF) grants (RS-2021-NR058898, RS-2023-00219517, and RS-2026-25488310) and the Main Research Program (E0210103) of the Korea Food Research Institute, all funded by the Korea government (MSIT), in addition to support from the National Institutes of Health (Grant ID: R01 DK129599 to Byung-Eun Kim).

## AUTHOR CONTRIBUTIONS

Y.-S.L. and H.-S.K. designed and performed experiments, analyzed and interpreted data, and wrote the manuscript; P.L.N. conducted bioinformatics analysis; J.L. and D.-I.K. generated *Ctr1*-AAV; J.L. and C.M. performed histological analysis; K.-O.C., B.-E.K., T.F.O., and J.A. analyzed and interpreted data and critically reviewed the manuscript; T.D. performed modeling; J.-S.K., C.H.J., and T.-I.J. conceived and designed the experiments and wrote the manuscript; All authors reviewed and edited the manuscript.

## REFERENCES

1. Tapiero, H., Townsend, D.M., and Tew, K.D. (2003). Trace elements in human physiology and pathology. Copper. Biomed. Pharmacother. 57, 386–398. 10.1016/s0753-3322(03)00012-x.

2. Ruiz, L.M., Libedinsky, A., and Elorza, A.A. (2021). Role of Copper on Mitochondrial Function and Metabolism. Front. Mol. Biosci. 8, 711227. 10.3389/fmolb.2021.711227.

3. Bost, M., Houdart, S., Oberli, M., Kalonji, E., Huneau, J.F., and Margaritis, I. (2016). Dietary copper and human health: Current evidence and unresolved issues. J. Trace Elem. Med. Biol. 35, 107–115. 10.1016/j.jtemb.2016.02.006.

4. Chun, H., Catterton, T., Kim, H., Lee, J., and Kim, B.E. (2017). Organ-specific regulation of ATP7A abundance is coordinated with systemic copper homeostasis. Sci Rep 7, 12001. 10.1038/s41598-017-11961-z.

5. Kim, B.-E., Nevitt, T., and Thiele, D.J. (2008). Mechanisms for copper acquisition, distribution and regulation. Nat. Chem. Biol. 4, 176–185. 10.1038/nchembio.72.

6. La Fontaine, S., and Mercer, J.F. (2007). Trafficking of the copper-ATPases, ATP7A and ATP7B: role in copper homeostasis. Arch Biochem Biophys 463, 149–167. 10.1016/j.abb.2007.04.021.

7. Liu, Y., Pilankatta, R., Hatori, Y., Lewis, D., and Inesi, G. (2010). Comparative features of copper ATPases ATP7A and ATP7B heterologously expressed in COS-1 cells. Biochemistry 49, 10006–10012. 10.1021/bi101423j.

8. Nose, Y., Kim, B.E., and Thiele, D.J. (2006). Ctr1 drives intestinal copper absorption and is essential for growth, iron metabolism, and neonatal cardiac function. Cell Metab. 4, 235–244. 10.1016/j.cmet.2006.08.009.

9. Kim, B.E., Turski, M.L., Nose, Y., Casad, M., Rockman, H.A., and Thiele, D.J. (2010). Cardiac copper deficiency activates a systemic signaling mechanism that communicates with the copper acquisition and storage organs. Cell Metab. 11, 353–363. 10.1016/j.cmet.2010.04.003.

10. Frontera, W.R., and Ochala, J. (2015). Skeletal muscle: a brief review of structure and function. Calcif. Tissue Int. 96, 183–195. 10.1007/s00223-014-9915-y.

11. Mukund, K., and Subramaniam, S. (2020). Skeletal muscle: A review of molecular structure and function, in health and disease. Wiley Interdiscip. Rev. Syst. Biol. Med. 12, e1462. 10.1002/wsbm.1462.

12. Schiaffino, S., and Reggiani, C. (2011). Fiber types in mammalian skeletal muscles. Physiol. Rev. 91, 1447–1531. 10.1152/physrev.00031.2010.

13. Pette, D., and Staron, R.S. (1997). Mammalian skeletal muscle fiber type transitions. Int. Rev. Cytol. 170, 143–223. 10.1016/s0074-7696(08)61622-8.

14. Song, M.Y., Han, C.Y., Moon, Y.J., Lee, J.H., Bae, E.J., and Park, B.H. (2022). Sirt6 reprograms myofibers to oxidative type through CREB-dependent Sox6 suppression. Nat. Commun. 13, 1808. 10.1038/s41467-022-29472-5.

15. Wen, W., Chen, X., Huang, Z., Chen, D., Yu, B., He, J., Zheng, P., Luo, Y., Yan, H., and Yu, J. (2021). Lycopene increases the proportion of slow-twitch muscle fiber by AMPK signaling to improve muscle anti-fatigue ability. J. Nutr. Biochem. 94, 108750. 10.1016/j.jnutbio.2021.108750.

16. Pereira, C.V., Peralta, S., Arguello, T., Bacman, S.R., Diaz, F., and Moraes, C.T. (2020). Myopathy reversion in mice after restauration of mitochondrial complex I. EMBO Mol. Med. 12, e10674. 10.15252/emmm.201910674.

17. Diaz, F., Thomas, C.K., Garcia, S., Hernandez, D., and Moraes, C.T. (2005). Mice lacking COX10 in skeletal muscle recapitulate the phenotype of progressive mitochondrial myopathies associated with cytochrome c oxidase deficiency. Hum. Mol. Genet. 14, 2737–2748. 10.1093/hmg/ddi307.

18. Tsukihara, T., Aoyama, H., Yamashita, E., Tomizaki, T., Yamaguchi, H., Shinzawa-Itoh, K., Nakashima, R., Yaono, R., and Yoshikawa, S. (1995). Structures of Metal Sites of Oxidized Bovine Heart Cytochrome c Oxidase at 2.8 Å. Science 269, 1069–1074. doi:10.1126/science.7652554.

19. Leary, S.C., Winge, D.R., and Cobine, P.A. (2009). “Pulling the plug” on cellular copper: the role of mitochondria in copper export. Biochim Biophys Acta 1793, 146–153. 10.1016/j.bbamcr.2008.05.002.

20. Bustos, R.I., Jensen, E.L., Ruiz, L.M., Rivera, S., Ruiz, S., Simon, F., Riedel, C., Ferrick, D., and Elorza, A.A. (2013). Copper deficiency alters cell bioenergetics and induces mitochondrial fusion through up-regulation of MFN2 and OPA1 in erythropoietic cells. Biochem. Biophys. Res. Commun. 437, 426–432. 10.1016/j.bbrc.2013.06.095.

21. Squitti, R., and Polimanti, R. (2013). Copper phenotype in Alzheimer’s disease: dissecting the pathway. Am J Neurodegener Dis 2, 46–56.

22. Fujisawa, C., Kodama, H., Sato, Y., Mimaki, M., Yagi, M., Awano, H., Matsuo, M., Shintaku, H., Yoshida, S., Takayanagi, M., et al. (2022). Early clinical signs and treatment of Menkes disease. Molecular Genetics and Metabolism Reports 31, 100849. 10.1016/j.ymgmr.2022.100849.

23. Vest, K.E., Paskavitz, A.L., Lee, J.B., and Padilla-Benavides, T. (2018). Dynamic changes in copper homeostasis and post-transcriptional regulation of Atp7a during myogenic differentiation. Metallomics 10, 309–322. 10.1039/c7mt00324b.

24. McCann, C., Quinteros, M., Adelugba, I., Morgada, M.N., Castelblanco, A.R., Davis, E.J., Lanzirotti, A., Hainer, S.J., Vila, A.J., Navea, J.G., and Padilla-Benavides, T. (2022). The mitochondrial Cu(+) transporter PiC2 (SLC25A3) is a target of MTF1 and contributes to the development of skeletal muscle in vitro. Front. Mol. Biosci. 9, 1037941. 10.3389/fmolb.2022.1037941.

25. Gregory, C.M., Vandenborne, K., and Dudley, G.A. (2001). Metabolic enzymes and phenotypic expression among human locomotor muscles. Muscle & Nerve 24, 387–393. 10.1002/1097-4598(200103)24:3<387::aid-mus1010>3.0.co;2-m.

26. Chennamsetty, I., Coronado, M., Contrepois, K., Keller, M.P., Carcamo-Orive, I., Sandin, J., Fajardo, G., Whittle, A.J., Fathzadeh, M., Snyder, M., et al. (2016). Nat1 Deficiency Is Associated with Mitochondrial Dysfunction and Exercise Intolerance in Mice. Cell Rep. 17, 527–540. 10.1016/j.celrep.2016.09.005.

27. Brischigliaro, M., and Zeviani, M. (2021). Cytochrome c oxidase deficiency. Biochim. Biophys. Acta Bioenerg. 1862, 148335. 10.1016/j.bbabio.2020.148335.

28. Furukawa, Y., Torres, A.S., and O’Halloran, T.V. (2004). Oxygenlinduced maturation of SOD1: a key role for disulfide formation by the copper chaperone CCS. EMBO J. 23, 2872–2881. 10.1038/sj.emboj.7600276.

29. Bertinato, J., and L’Abbé, M.R. (2003). Copper modulates the degradation of copper chaperone for Cu,Zn superoxide dismutase by the 26 S proteosome. J. Biol. Chem. 278, 35071–35078. 10.1074/jbc.M302242200.

30. Ryzhkova, A.I., Sazonova, M.A., Sinyov, V.V., Galitsyna, E.V., Chicheva, M.M., Melnichenko, A.A., Grechko, A.V., Postnov, A.Y., Orekhov, A.N., and Shkurat, T.P. (2018). Mitochondrial diseases caused by mtDNA mutations: a mini-review. Ther. Clin. Risk Manag. 14, 1933–1942. 10.2147/tcrm.S154863.

31. Ahmed, S.T., Craven, L., Russell, O.M., Turnbull, D.M., and Vincent, A.E. (2018). Diagnosis and Treatment of Mitochondrial Myopathies. Neurotherapeutics 15, 943–953. 10.1007/s13311-018-00674-4.

32. Aptekmann, A.A., Buongiorno, J., Giovannelli, D., Glamoclija, M., Ferreiro, D.U., and Bromberg, Y. (2022). mebipred: identifying metal-binding potential in protein sequence. Bioinformatics 38, 3532–3540. 10.1093/bioinformatics/btac358.

33. Goldman, A., Mullokandov, M., Zaltsman, Y., Regev, L., Levin-Zaidman, S., and Gross, A. (2024). MTCH2 cooperates with MFN2 and lysophosphatidic acid synthesis to sustain mitochondrial fusion. EMBO Rep. 25, 45–67. 10.1038/s44319-023-00009-1.

34. Grinberg, M., Schwarz, M., Zaltsman, Y., Eini, T., Niv, H., Pietrokovski, S., and Gross, A. (2005). Mitochondrial carrier homolog 2 is a target of tBID in cells signaled to die by tumor necrosis factor alpha. Mol Cell Biol 25, 4579–4590. 10.1128/mcb.25.11.4579-4590.2005.

35. Buzaglo-Azriel, L., Kuperman, Y., Tsoory, M., Zaltsman, Y., Shachnai, L., Zaidman, S.L., Bassat, E., Michailovici, I., Sarver, A., Tzahor, E., et al. (2017). Loss of Muscle MTCH2 Increases Whole-Body Energy Utilization and Protects from Diet-Induced Obesity. Cell Rep. 18, 1335–1336. 10.1016/j.celrep.2017.01.046.

36. Guna, A., Stevens, T.A., Inglis, A.J., Replogle, J.M., Esantsi, T.K., Muthukumar, G., Shaffer, K.C.L., Wang, M.L., Pogson, A.N., Jones, J.J., et al. (2022). MTCH2 is a mitochondrial outer membrane protein insertase. Science 378, 317–322. 10.1126/science.add1856.

37. Kang, Y., and Chen, L. (2023). Structural basis for the binding of DNP and purine nucleotides onto UCP1. Nature 620, 226–231. 10.1038/s41586-023-06332-w.

38. Pebay-Peyroula, E., Dahout-Gonzalez, C., Kahn, R., Trézéguet, V., Lauquin, G.J.M., and Brandolin, G. (2003). Structure of mitochondrial ADP/ATP carrier in complex with carboxyatractyloside. Nature 426, 39–44. 10.1038/nature02056.

39. Waterhouse, A., Bertoni, M., Bienert, S., Studer, G., Tauriello, G., Gumienny, R., Heer, F.T., de Beer, T.A.P., Rempfer, C., Bordoli, L., et al. (2018). SWISS-MODEL: homology modelling of protein structures and complexes. Nucleic Acids Res. 46, W296–w303. 10.1093/nar/gky427.

40. Forrest, I., Conway, L.P., Jadhav, A.M., Gathmann, C., Chiu, T.-Y., Chaheine, C.M., Estrada, M., Shrestha, A., Reitsma, J.M., Warder, S.E., et al. (2025). Proteome-Wide Discovery of Degradable Proteins Using Bifunctional Molecules. bioRxiv, 2025.2003.2021.644652. 10.1101/2025.03.21.644652.

41. Bahat, A., Goldman, A., Zaltsman, Y., Khan, D.H., Halperin, C., Amzallag, E., Krupalnik, V., Mullokandov, M., Silberman, A., Erez, A., et al. (2018). MTCH2-mediated mitochondrial fusion drives exit from naïve pluripotency in embryonic stem cells. Nat. Commun. 9, 5132. 10.1038/s41467-018-07519-w.

42. Yoneshiro, T., Wang, Q., Tajima, K., Matsushita, M., Maki, H., Igarashi, K., Dai, Z., White, P.J., McGarrah, R.W., Ilkayeva, O.R., et al. (2019). BCAA catabolism in brown fat controls energy homeostasis through SLC25A44. Nature 572, 614–619. 10.1038/s41586-019-1503-x.

43. Verkerke, A.R.P., Shi, X., Li, M., Higuchi, Y., Yamamuro, T., Katoh, D., Nishida, H., Auger, C., Abe, I., Gerszten, R.E., and Kajimura, S. (2024). SLC25A48 controls mitochondrial choline import and metabolism. Cell Metabolism 36, 2156–2166.e2159. 10.1016/j.cmet.2024.07.010.

44. Nose, Y., Wood, L.K., Kim, B.-E., Prohaska, J.R., Fry, R.S., Spears, J.W., and Thiele, D.J. (2010). Ctr1 Is an Apical Copper Transporter in Mammalian Intestinal Epithelial Cells in Vivo That Is Controlled at the Level of Protein Stability*. J. Biol. Chem. 285, 32385–32392. 10.1074/jbc.M110.143826.

45. Soma, S., Latimer, A.J., Chun, H., Vicary, A.C., Timbalia, S.A., Boulet, A., Rahn, J.J., Chan, S.S.L., Leary, S.C., Kim, B.E., et al. (2018). Elesclomol restores mitochondrial function in genetic models of copper deficiency. Proc. Natl Acad. Sci. USA 115, 8161–8166. 10.1073/pnas.1806296115.

46. Yuan, S., Korolnek, T., and Kim, B.E. (2022). Oral Elesclomol Treatment Alleviates Copper Deficiency in Animal Models. Front. Cell Dev. Biol. 10, 856300. 10.3389/fcell.2022.856300.

47. Guthrie, L.M., Soma, S., Yuan, S., Silva, A., Zulkifli, M., Snavely, T.C., Greene, H.F., Nunez, E., Lynch, B., De Ville, C., et al. (2020). Elesclomol alleviates Menkes pathology and mortality by escorting Cu to cuproenzymes in mice. Science 368, 620–625. 10.1126/science.aaz8899.

48. Hood, D.A., Memme, J.M., Oliveira, A.N., and Triolo, M. (2019). Maintenance of Skeletal Muscle Mitochondria in Health, Exercise, and Aging. Annu. Rev. Physiol. 81, 19–41. 10.1146/annurev-physiol-020518-114310.

49. Cobine, P.A., Moore, S.A., and Leary, S.C. (2021). Getting out what you put in: Copper in mitochondria and its impacts on human disease. Biochim. Biophys. Acta Mol. Cell Res. 1868, 118867. 10.1016/j.bbamcr.2020.118867.

50. Boulet, A., Vest, K.E., Maynard, M.K., Gammon, M.G., Russell, A.C., Mathews, A.T., Cole, S.E., Zhu, X., Phillips, C.B., Kwong, J.Q., et al. (2018). The mammalian phosphate carrier SLC25A3 is a mitochondrial copper transporter required for cytochrome c oxidase biogenesis. J. Biol. Chem. 293, 1887–1896. 10.1074/jbc.RA117.000265.

51. Xu, W., Barrientos, T., and Andrews, N.C. (2013). Iron and copper in mitochondrial diseases. Cell Metab. 17, 319–328. 10.1016/j.cmet.2013.02.004.

52. Murata, D., Roy, S., Lutsenko, S., Iijima, M., and Sesaki, H. (2024). Slc25a3-dependent copper transport controls flickering-induced Opa1 processing for mitochondrial safeguard. Developmental Cell 59, 2578–2592.e2577. 10.1016/j.devcel.2024.06.008.

53. Chourasia, S., Petucci, C., Shoffler, C., Abbasian, D., Wang, H., Han, X., Sivan, E., Brandis, A., Mehlman, T., Malitsky, S., et al. (2025). MTCH2 controls energy demand and expenditure to fuel anabolism during adipogenesis. EMBO J. 44, 1007–1038. 10.1038/s44318-024-00335-7.

54. Grevendonk, L., Connell, N.J., McCrum, C., Fealy, C.E., Bilet, L., Bruls, Y.M.H., Mevenkamp, J., Schrauwen-Hinderling, V.B., Jörgensen, J.A., Moonen-Kornips, E., et al. (2021). Impact of aging and exercise on skeletal muscle mitochondrial capacity, energy metabolism, and physical function. Nat. Commun. 12, 4773. 10.1038/s41467-021-24956-2.

55. Allodi, I., Montañana-Rosell, R., Selvan, R., Löw, P., and Kiehn, O. (2021). Locomotor deficits in a mouse model of ALS are paralleled by loss of V1-interneuron connections onto fast motor neurons. Nat. Commun. 12, 3251. 10.1038/s41467-021-23224-7.

56. Białobrodzka, E., Flis, D.J., Akdogan, B., Borkowska, A., Wieckowski, M.R., Antosiewicz, J., Zischka, H., Dzik, K.P., Kaczor, J.J., and Ziolkowski, W. (2024). Amyotrophic Lateral Sclerosis and swim training affect copper metabolism in skeletal muscle in a mouse model of disease. Muscle & Nerve 70, 1111–1118. 10.1002/mus.28237.

57. Ogbonmide, T., Rathore, R., Rangrej, S.B., Hutchinson, S., Lewis, M., Ojilere, S., Carvalho, V., and Kelly, I. (2023). Gene Therapy for Spinal Muscular Atrophy (SMA): A Review of Current Challenges and Safety Considerations for Onasemnogene Abeparvovec (Zolgensma). Cureus 15, e36197. 10.7759/cureus.36197.

58. Mendell, J.R., Al-Zaidy, S.A., Rodino-Klapac, L.R., Goodspeed, K., Gray, S.J., Kay, C.N., Boye, S.L., Boye, S.E., George, L.A., Salabarria, S., et al. (2021). Current Clinical Applications of In Vivo Gene Therapy with AAVs. Mol. Ther. 29, 464–488. 10.1016/j.ymthe.2020.12.007.

59. Mah, C.S., Soustek, M.S., Todd, A.G., McCall, A., Smith, B.K., Corti, M., Falk, D.J., and Byrne, B.J. (2013). Adeno-associated virus-mediated gene therapy for metabolic myopathy. Hum. Gene. Ther. 24, 928–936. 10.1089/hum.2013.2514.

