## Supplementary Data for "Copper deficiency drives OXPHOS impairment and mitochondrial hyperfusion via MTCH2 in skeletal muscle"

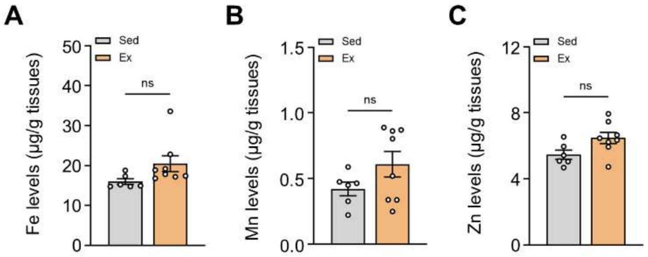


**Figure S1. Exercise does not alter iron, manganese, or zinc content in skeletal muscle.** (A-C) Quantification of (A) iron (Fe), (B) manganese (Mn), and (C) zinc (Zn) levels by ICP-MS in the gastrocnemius (GA) skeletal muscles of sedentary and exercise-trained wild-type mice (*n* = 6-8). Data are presented as mean ± SEM. Statistical differences were determined by two-tailed Student's t-test.

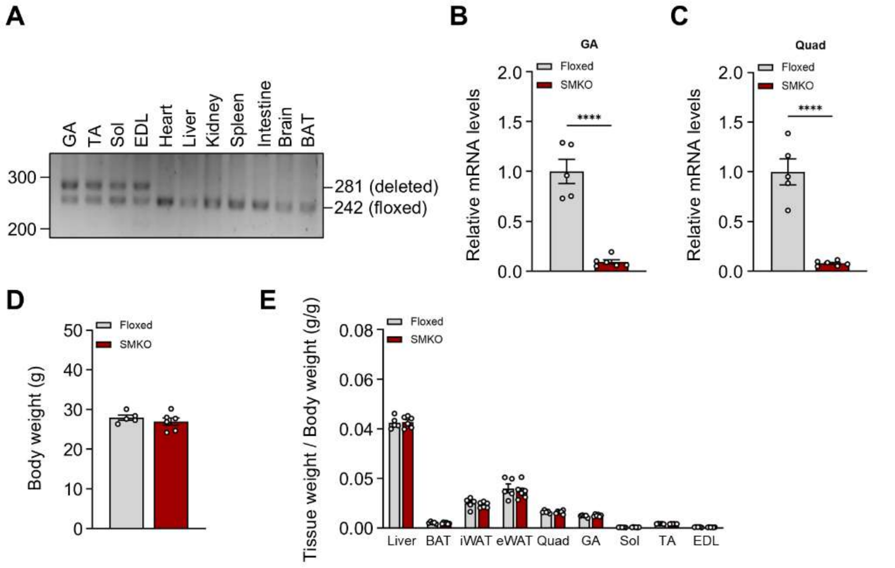


**Figure S2. Validation of skeletal muscle-specific *Ctr1* deletion and assessment of body and tissue wights.** (A) Analysis of *Ctr1* genomic excision across various tissues in SMKO mice. (B and C) Relative *Ctr1* mRNA expression in the (B) GA and (C) Quadriceps (Quad) muscles of Floxed and SMKO mice ($n$ = 5-6). (D and E) (D) Total body weight and (E) isolated tissue weights normalized to body weight of Floxed and SMKO mice ($n$ = 5-6). Data are presented as mean ± SEM. Statistical differences were determined by two-tailed Student's t-test.

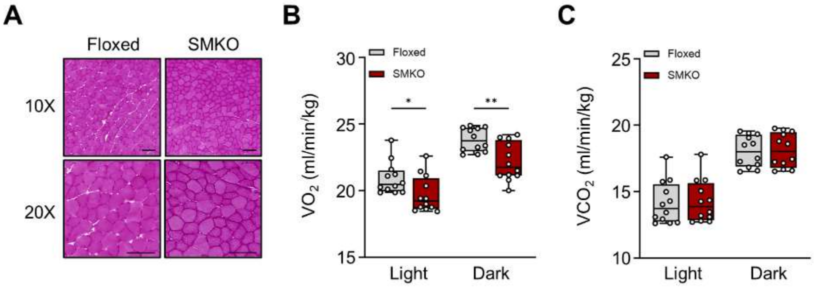


**Figure S3. SMKO mice exhibit reduced systemic oxygen consumption without gross histological alterations.** (A) Representative H&E staining of GA muscles from Floxed and SMKO mice. Scale bar = 100 µm. (B and C) Whole-body indirect calorimetry analysis evaluating (B) oxygen consumption ($VO_2$) and (C) carbon dioxide production ($VCO_2$) in Floxed and SMKO mice. Data are presented as mean ± SEM. Statistical differences were determined by two-way ANOVA with Šídák's multiple comparisons test.

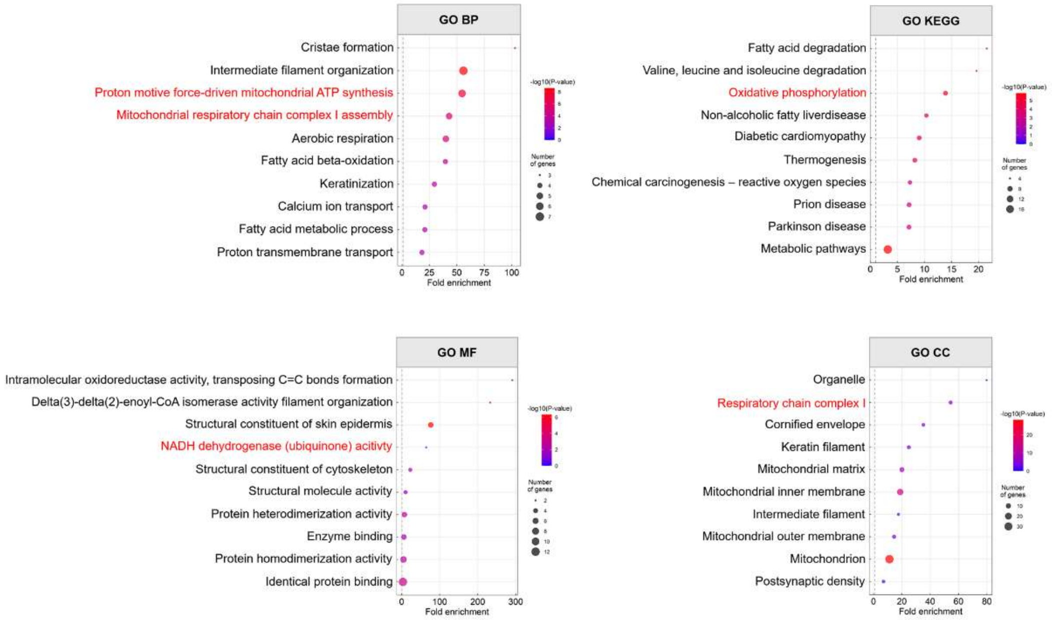


**Figure S4. Upregulated proteins in SMKO muscle are enriched for Complex I and ATP synthesis pathway.** Gene Ontology (GO) enrichment analysis of significantly upregulated proteins in the GA muscle of SMKO mice compared to Floxed controls.

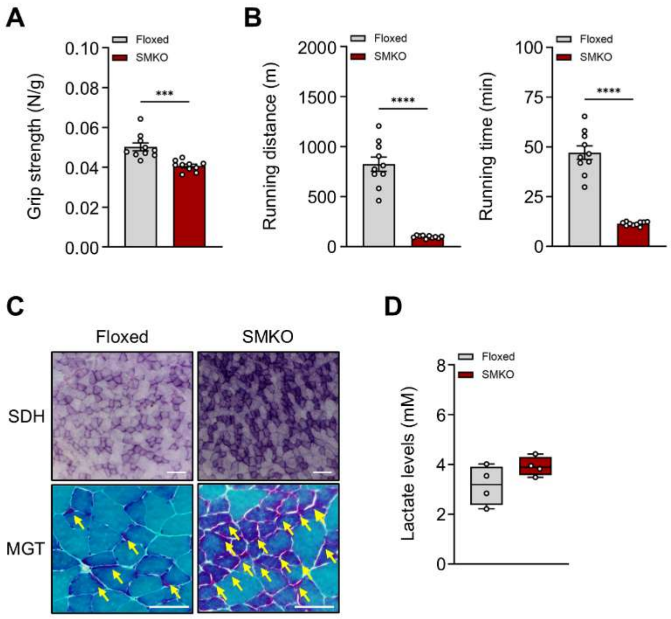


**Figure S5. Female SMKO mice exhibit comparable hallmarks of mitochondrial myopathy.** (A) Forelimb grip strength of Floxed and SMKO female mice (*n* = 10). (B) Total running distance and time to exhaustion during treadmill endurance testing (*n* = 10). (C) Histological analysis in GA muscles from Floxed and SMKO female mice. Succinate dehydrogenase (SDH) and Modified Gomori trichorme (MGT) staining. Scale bar = 100 µm (SDH) or 50 µm (MGT). (D) Blood lactate concentration in Floxed and SMKO female mice (*n* = 4). Data are presented as mean ± SEM. Statistical differences were determined by two-tailed Student's t-test.

**A**

*SMKO vs Floxed*

| Gene symbol | Protein | Accession | SMKO vs | Ions with binding potential |
|---|---|---|---|---|
| Eci1 | Enoyl-CoA delta isomerase 1, mitochondrial | P42125 | 2.012 | $Cu^1$, $Zn^1$, $Ca^1$ |
| Mtch2 | Mitochondrial carrier homolog 2 | Q791V5 | 1.852 | $Cu^1$ |
| Ttn | Isoform 3 of Titin | A2ASS6-3 | 0.506 | $Cu^1$,$Fe^1$,$K^1$,$Mg^1$,$Mn^1$,$Na^1$,$Ni^1$,$Zn^1$,$Ca^1$ |
| Nap1l4 | Nap1l4 protein | B7ZNL2 | 0.435 | $Cu^1$,$Ca^1$ |
| Mtco2 | Cytochrome c oxidase subunit 2 | P00405 | 0.064 | $Cu^1$,$Mg^1$,$Ni^1$,$Zn^1$,$Co^1$ |
| Mtco1 | Cytochrome c oxidase subunit 1 | P00397 | 0.056 | $Cu^1$,$Fe^1$,$Mg^1$,$Na^1$,$Ni^1$ |

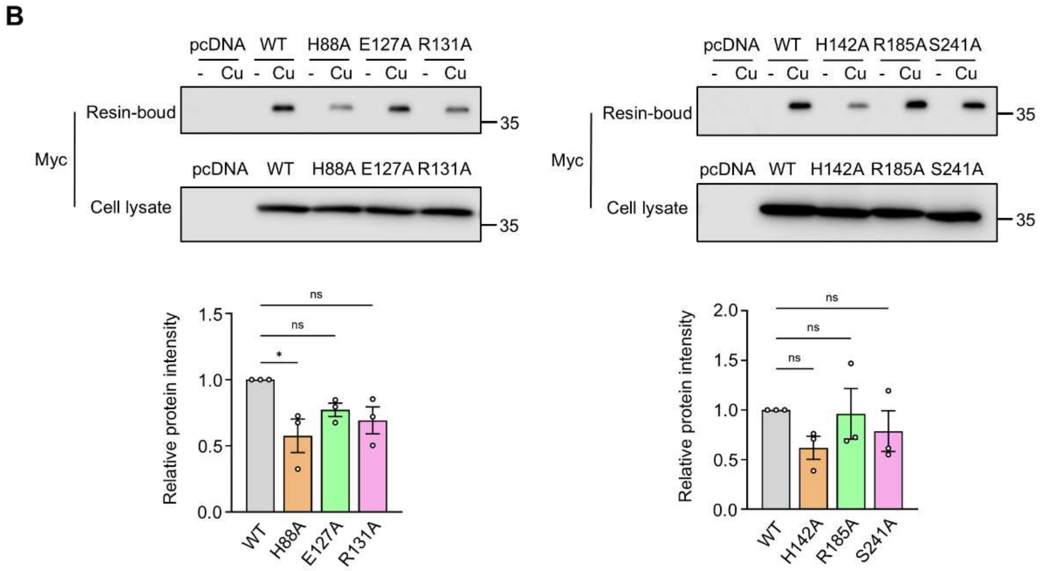


**Figure S6. MTCH2 is upregulated under copper deficiency, and its copper coordination is independent of internal nitrogen- and oxygen-rich residues.** (A) Proteomic upregulation of MTCH2 in SMKO muscle and computational prediction of its copper-binding potential utilizing the MeBiPred analysis tool. (B) *In vitro* copper-binding assay utilizing HEK293T cells expressing wild-type (WT) or internal nitrogen/oxygen-rich alanine-substitution mutants of pCMV3-MTCH2-Myc. Bound protein fractions were normalized to total Myc expression in whole-cell lysates. $n = 3$ independent experiments. Data are presented as mean ± SEM. Statistical differences were determined by two-way ANOVA with Šídák's multiple comparisons test.

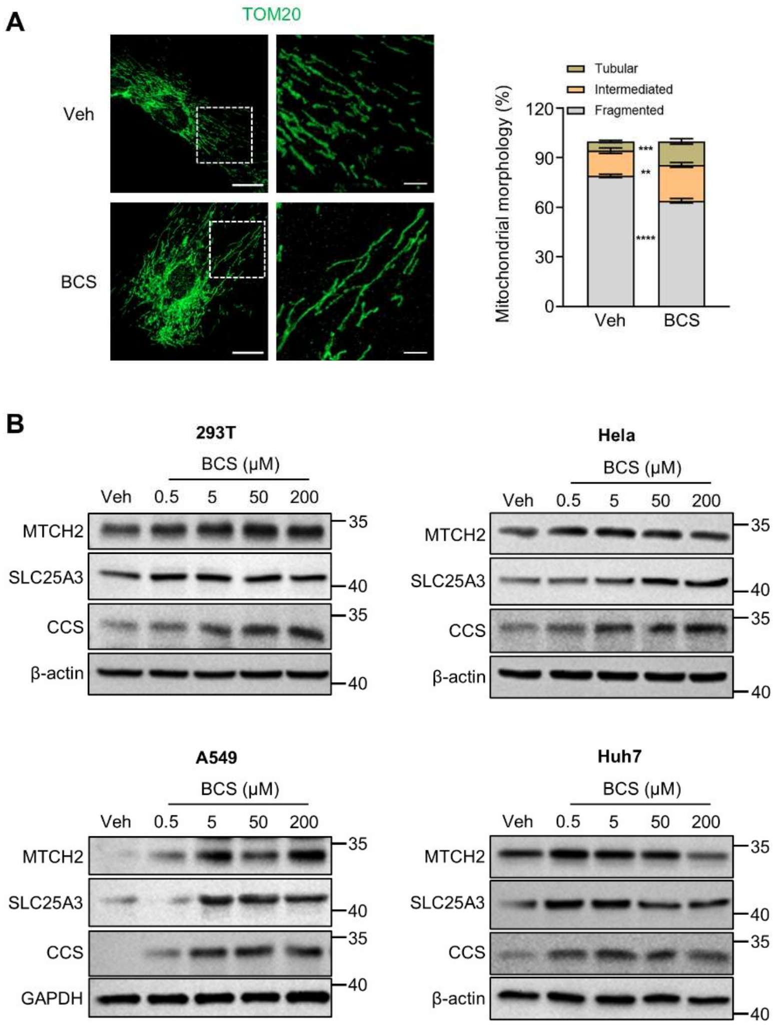


**Figure S7. Copper deficiency induces MTCH2-dependent mitochondrial hyperfusion.** (A) Immunofluorescence imaging of mitochondrial morphology in human skeletal muscle myoblast (HSMM) treated with BCS (50 μM). Scale bars = 20 μm (main) and 5 μm (inset). Mitochondrial network morphology was quantified using ImageJ. (B) Immunoblot analysis of MTCH2 and SLC25A3 expression across various specified cell lines treated with vehicle or BCS for 24 h. Data are presented as mean ± SEM. Statistical differences were determined by two-way ANOVA with Tukey's test.

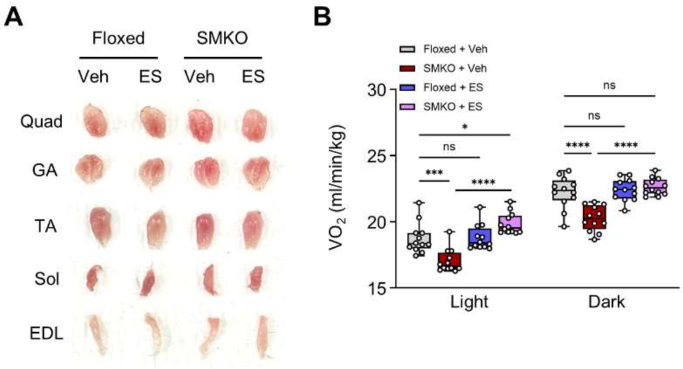


**Figure S8. Elesclomol normalizes macroscopic tissue color and systemic oxygen consumption in SMKO mice.** Elesclomol (ES) was administrated subcutaneously to Floxed and SMKO mice for 2 weeks. (A) Representative gross macroscopic images of skeletal muscles from vehicle- and ES-treated mice. (B) Whole-body oxygen consumption rate ($VO_2$) ($n$ = 5-7). Data are presented as mean ± SEM. Statistical differences were determined by two-way ANOVA with Tukey's test.

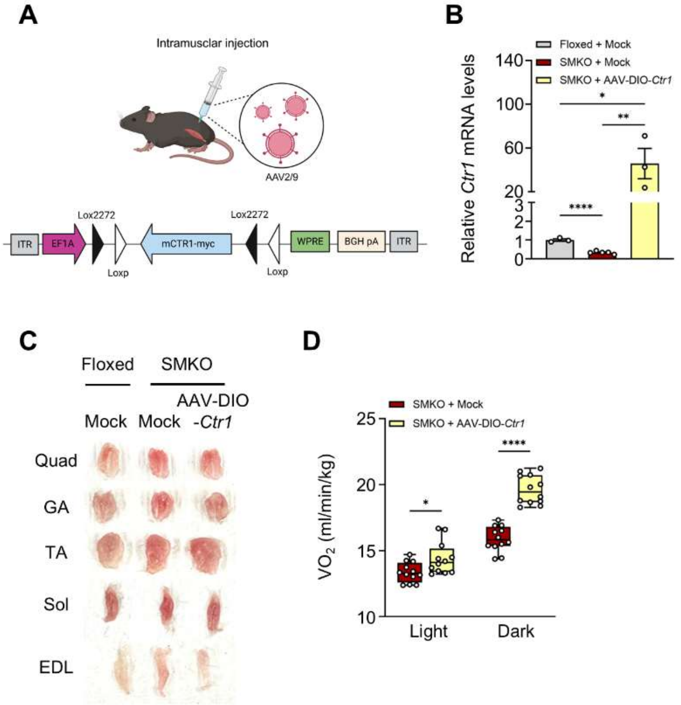


**Figure S9. AAV-DIO-mediated *Ctr1* overexpression restores myofiber composition and oxygen consumption in SMKO mice.** A single intramuscular injection of AAV-DIO-*Ctr1* was administered to SMKO mice, and assessments were performed after 5 weeks. (A) Schematic representations of the genetic configurations for the AAV-DIO-*Ctr1* vector. (B) Relative mRNA expression of Ctr1 ($n$ = 3-5). (C) Representative macroscopic images of skeletal muscles showing tissue color. (D) Whole-body oxygen uptake ($VO_2$) ($n$ = 3). Data are presented as mean ± SEM. Statistical differences were determined by two-tailed Student's t-test (B) and two-way ANOVA with Šídák's multiple comparisons test (D).
